# A neural turning point - the EEG P3 component tracks unfolding changes of mind

**DOI:** 10.1101/2020.11.18.388363

**Authors:** Elisabeth Parés-Pujolràs, Jeremy Hatchuel, Patrick Haggard

## Abstract

The ability to change one’s mind is a key feature of human cognition. Yet, the neural mechanisms underpinning our capacity to change our minds remain poorly understood. Here, we investigated the neural correlates of evidence accumulation and changes of mind in a two-step sequential sampling task. Participants provided a first, quick guess regarding the relative frequencies of target letters in a visual stream, followed by a slower, more deliberate decision. We found that the P3 amplitude evoked by successive target letters tracks an internal signed decision variable and predicts choices on a single-trial level. Moreover, this neural decision variable offers new insights into the dynamics of changes of mind. In particular, we show that the start of evidence accumulation after the early decision constitutes a neural turning point: the P3 evoked by the first letter contrary to the initial decision can be used to predict subsequent changes of mind. Our results highlight a critical interaction between the processing of external evidence and endogenous modulations of decisional parameters that facilitate reversing an original decision.

## 1. Introduction

Voluntary control of behaviour requires the ability to dynamically integrate internal states and external evidence to successfully achieve one’s goals. The ability to commit to an initial choice and persevere on a given action path enables agents to maintain and fulfil intentions over long time spans. Yet, excessive perseverance can turn into maladaptive stubbornness if agents are unable to dynamically adapt their behaviour and change their minds if necessary ^1^. How do people decide whether to stick to their initial choices or rather change their minds?

Accumulation-to-bound theories of decision-making suggest that evidence is continuously sampled and integrated in an internal decision variable until a threshold is reached, triggering the corresponding action^2,3^. Importantly, most decisions are not irreversible, and evidence accumulation typically continues even after an action has been executed. In simple perceptual decision-making tasks, changes of mind typically improve accuracy ^4–6^, and thus provide an important mechanism for behavioural optimization ^7,8^. Computationally, changes of mind can be modelled as a reversal in the sign of the decision variable, which drifts away from the bound hit at the time of the first decision and moves closer towards the opposite bound^9^. While neural signals that correspond to such changes have been identified in non-human primates^5^, identifying reliable markers of unfolding changes of mind in humans remains a challenge. As a result, understanding of the cognitive processes involved remains correspondingly limited.

Importantly, external input is not the only factor determining the evolution of such variables. For example, different contexts may require agents to make fast decisions, while others may call for accurate choices, perhaps at the cost of slower action^10^. Agents are able to tune their decision-making processes at will to reflect their current prioritisation of speed or accuracy, and these modulations result in distinct changes in behaviour with identifiable neural correlates^11,12^. Further, it has been suggested that in any given decision an internal growing urgency signal modulates the impact of evidence on decision variables ^11–14^, pushing the decision variable closer to threshold as time elapses to prevent agents from spending too much time deliberating.

Moreover, after a decision has been made, confidence in initial choices can further modulate how new information is processed ^15–17^. High confidence choices are followed by modulations of the starting point and gain in computational models of post-decisional evidence accumulation that can account for the well-known confirmation bias ^16^. Conversely, low confidence in initial choices leads to an increased probability of changing one’s mind ^16^.

In this study, we sought to identify a neural marker that would allow us to track unfolding changes of mind as they unfold, and to investigate the interaction between endogenous factors and the processing of external evidence that underlie flexible decision-making. Recent research has shown that the classic P3 of the human EEG, and the related Centro Parietal Positivity (CPP), encode a build-to-threshold variable that tracks decision-making processes in real-time ^18–20^. These studies have provided key insights into the neural correlates of evidence accumulation in humans. However, only recently have such signals been used to track how alternatives compete during choice formation, and *which* decision participants will eventually make ^21^. Here, we build on this research to test whether the P3 can be further used to track unfolding changes of mind.

In our task, participants viewed a continuous stream of letters. Their task was to decide which of two categories of stimuli (*b* and *d* vs. *p* and *q*) were presented more frequently. They were instructed to make a fast guess early in the trial. The stimuli presentation continued after their initial guess, and they were asked to make a second action reporting their final decision without any strong time pressure. They thus had a chance to revise their initial decision and change their mind. We predicted that, if the P3 indeed tracks the state of an internal decision variable, the difference in P3 amplitudes evoked by the two sets of target letters should reflect the participants’ choices and track changes of mind as they unfold in our continuous task. Our use of discrete, sequential evidence items further allowed us to test whether endogenous modulations of neural responses to confirmatory and disconfirmatory evidence following initial choices facilitated subsequent changes of mind.

## 2. Results

### 2.1. Task & design

Participants (n = 19) performed a simple sequential sampling decision-making task (*Figure 1)*. Each trial started with a grey background, and a 3.75Hz letter stream containing relevant and distractor stimuli was presented screen. There were two sets of task-relevant letters, to which a task-relevant colour corresponded. Thus ‘b’ and ‘d’ corresponded to the colour blue, and ‘p’ and ‘q’ to pink. Participants were instructed to monitor the letter stream and decide which set of targets was appearing more frequently (i.e. *bd* or *pq*). The participants’ task was to make sure that the background colour of the screen matched the more frequent group of letters (i.e. if the most frequent set of targets was *bd*, the screen should be blue. If the most frequent set of targets was *pq*, the screen should be pink). Participants were asked to execute two actions to indicate their decisions: a fast and a slow one. The first decision (Action 1) had to be as quick as possible, since their potential reward decreased up to the time of their first action. However, it had to be evidence-informed to a certain extent because changes of mind to correct an initial error were penalised (see *Methods*). The second decision (Action 2) had to be accurate, since if they made the wrong choice they would lose money. Trial duration was always the same, regardless of the timing of their actions: participants always had to wait for all of the 100 letters to be presented before the trial ended. In some trials, participants chose the same colour both for the first and the second action (*No-CoM,* no Change of Mind). In some other trials, participants chose different colours in Action 1 and Action 2 (*CoM,* Change of Mind). At the end of each trial, participants were asked to estimate the time of their first and second actions on two visual analogue scales (VAS), and to report how confident they were on their final (i.e. second) decision on a further VAS.

**Figure 1.**
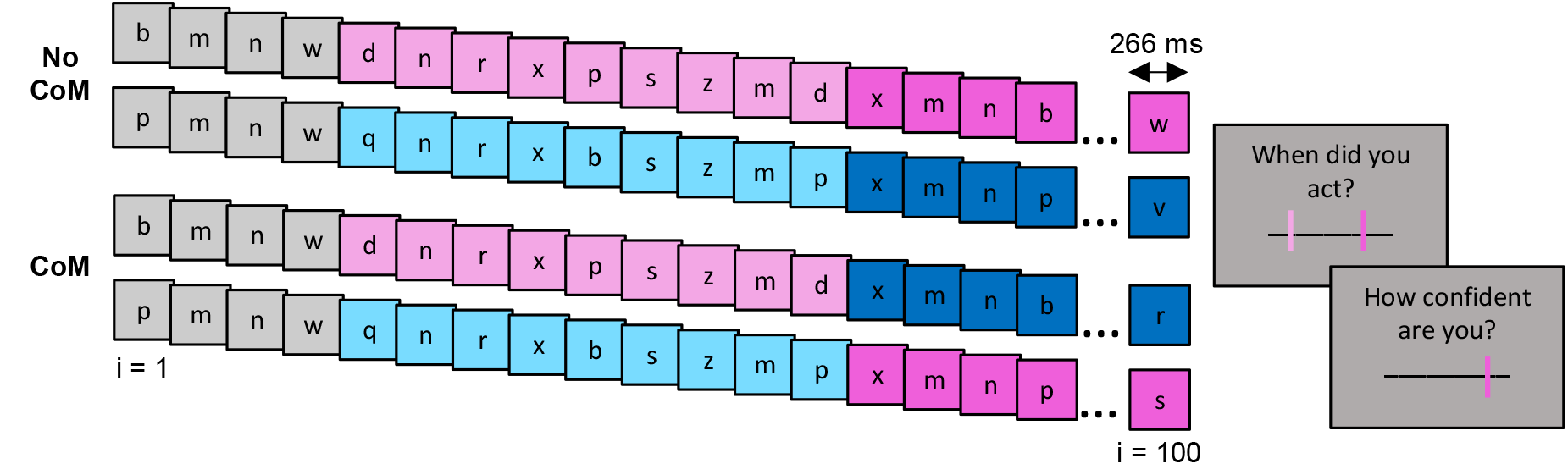
Task structure. Each trial started with a grey background. A letter stream of 100 letters was presented for 26.6 seconds, and participants had to decide which of the two target letter sets was most frequent (blue targets: ‘b’ and ‘d’, or pink targets: ‘p’ and ‘q’). Their task was to make sure that the colour of the screen matched the most frequent set of targets (e.g. if ‘p’ and ‘q’ were most frequent, the screen should be pink). They were required to execute two actions. Action 1 had to be as fast as possible, since their maximal reward decreased until the time of the first action. Action 2 had to be as accurate as possible. If they made a mistake, they lost money. Participants sometimes chose the same colour both in Action 1 and Action 2 (*No CoM,* no change of mind), but sometimes they changed of mind (*CoM*). At the end of the trial, they were asked to provide an estimate of the time at which they executed each action, and a separate judgement about their confidence in their final decision (Action 2).

There were two experimental conditions randomised within blocks. In the *Informative* condition (one third of trials, n = 38), the most frequent set of letters would appear twice as frequently (12.5% of total letters in the stream) as the less frequent set (6.25% of letters). Thus, the perceptual decision was easy. Half of these informative trials required the keypress indicating Blue (p(*bd*) > p(*pq*)) and the other half Pink (p(*bd*) < p(*pq*)). In the *Neutral* condition (two thirds of trials, n = 76), both sets of target letters were presented at the same frequency. Thus, overall, there was no strong evidence for or against any option in any given trial. On average, 16% of all letters presented were *bd* or *pq* targets. More neutral trials than informative ones were included in the design to maximise the number of trials in which participants would change their mind. Participants were not informed of the existence of neutral trials. Target letters were presented randomly but were always separated by at least 2 distractors (0.532 s interval).

Participants performed the task well. On average, they executed two actions in >90% of trials across conditions (Missing Action 2: *M =* 5.62%, *SD =* 3.69). Trials with missing responses were excluded from further analysis. In the informative conditions, their final choice was accurate on 94.03% (*SD* = 5.80%) of trials. Incorrect trials were not analysed further. Participants changed their mind significantly (*t*_(18)_ = 4.11, *p <* 0.001) less often in the Informative condition (*M* = 23.77%, *SD =* 7.92%) than in the Neutral condition (*M =* 35.87%*, SD=*12.93%). See *Supplementary Note 1* for full details of the behavioural results.

For analysis purposes, we grouped together the trials where participants chose Blue and those where they chose Pink, and we refer to the evidence as “response-congruent” or “response-incongruent” with respect to each action. That is, if participants chose ‘blue’, ‘b’ and ‘d’ preceding that action are response-congruent evidence, whereas ‘p’ and ‘q’ are labelled response-incongruent evidence. Further, when specifically referring to Action 2, we refer to evidence as “confirmatory” if it is congruent with the first response (e.g. *bd* if participants chose ‘Blue’ in Action 1), and “disconfirmatory” if it is incongruent with the first response (e.g. *pq* if participants chose Blue in Action 1).

### 2.2. The P3 encodes unfolding choices both in CoM and No-CoM trials

To study the neural correlates of evidence accumulation during our task, we investigated the evolution of the P3 amplitude evoked by sequential stimuli over the course of the trial. Each trial could be conceptually divided in two evidence accumulation processes: a first one from trial start leading to Action 1, and a second, slow one from Action 1 to Action 2. We interpret the P3 amplitudes evoked by the first piece of evidence presented in each decision-making process as a marker of the starting point for such decision (termed *S1* and *S2* for Action 1 and Action 2, respectively). Similarly, we interpret the P3 amplitudes evoked by the last piece of evidence as a proxy for the end-point of the decision process, or the decision threshold (*E1* and *E2* for Action 1 and Action 2, respectively; see *Figure 2a,d*).

**Figure 2.**
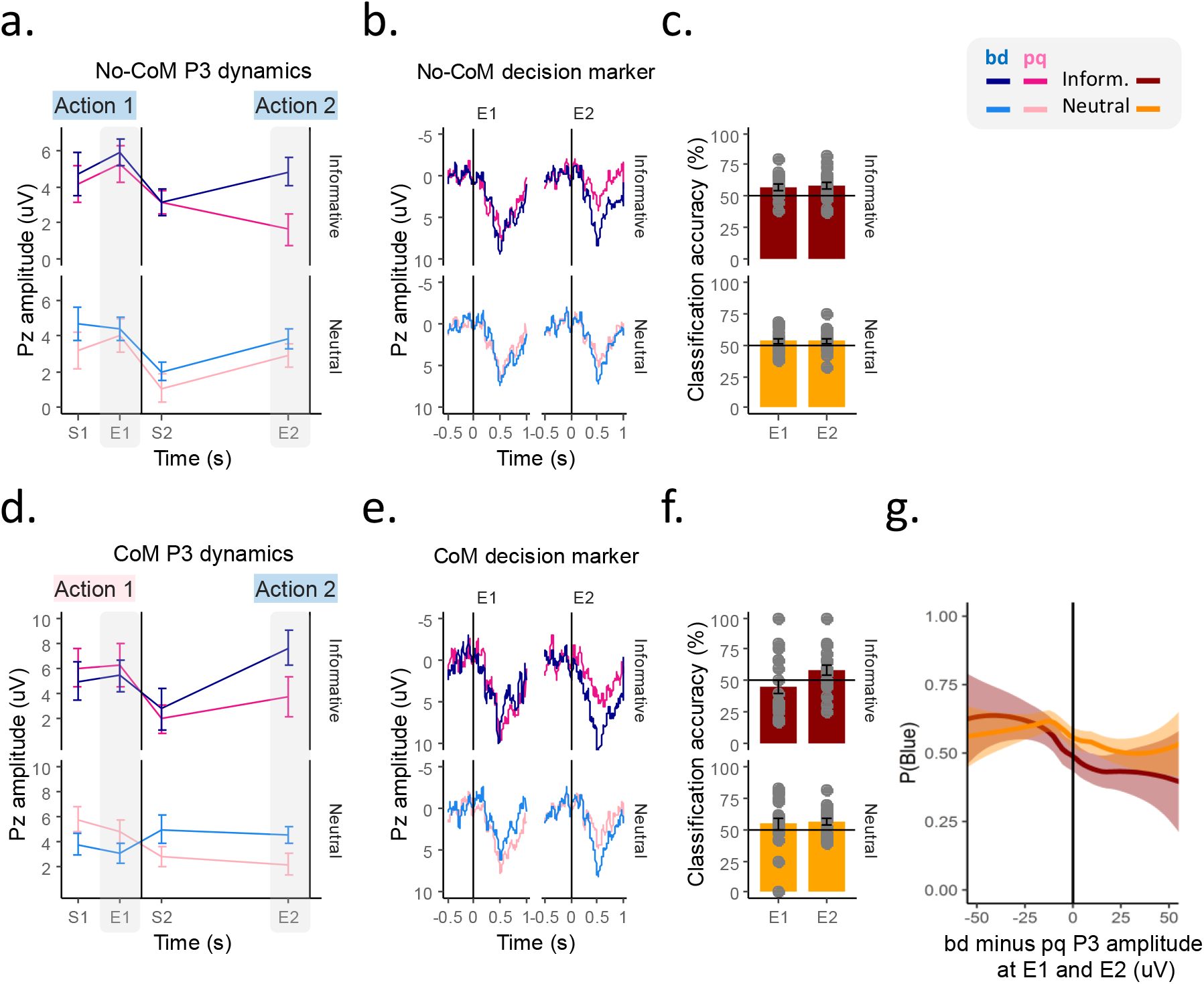
Dynamic changes in the P3 amplitude encode unfolding changes of mind. Trials in which participants chose the same colour in both Action 1 and Action 2 are illustrated on the top row (No-CoM), and those where they changed their mind in the bottom row (CoM). The graphs describe the full dataset. For illustration purposes, No-CoM trials (***a-c***) are color-coded as if participants had always chosen ‘Blue’. Similarly, CoM trials (***d-f***) are illustrated as if participants had always chosen ‘Pink’ at Action 1 and ‘Blue’ at Action 2. ***a,d.*** Line graphs illustrate the grand averaged (+SEM) amplitude of the P3 evoked by the first (*S1, S2)* and last (*E1, E2)* piece of evidence presented during the decision-making processes leading to Action 1 or Action 2, for both types of evidence. ***b, e.*** Grand-averaged ERP traces at Pz locked to the last piece of evidence presented before Action 1 (*E1*) and before Action 2 (*E2*). When participants did not change their mind, response-congruent evidence evoked higher P3 amplitudes compared to response-incongruent evidence before both actions (***b***). When participants changed their mind, the dominant P3 amplitudes numerically reversed between Action 1 and Action 2 (***e***). ***c,f.*** Mean (+SEM) choice classification accuracy of a P3-based heuristic. Overimposed dots illustrate individual participant means. The signed difference in P3 amplitudes evoked by competing evidence types at *E1* and *E2* could be used to significantly decode participants’ choices in both actions in the Neutral condition (both in CoM and No-CoM trials), but only Action 2 choices in the Informative CoM trials (***f***). ***g.*** Mean (+SE) probability of choosing ‘Blue’ (y = 1) or ‘Pink’ (y = 0) as a function of the difference in the P3 evoked by the last pieces of evidence before action, pooled across all trial types (CoM/No-CoM) and actions (Action 1/Action 2).

To investigate whether the P3 tracked categorical decisions and changes of mind, we compared the last amplitudes of the P3 evoked by of different types of evidence (*bd/pq*) before each action (*E1/E2*), in all conditions (Informative/Neutral). We focussed on the P3 components evoked by the last pieces of evidence presented before action under the assumption that neural markers of choice are strongest towards the end of decision-making processes^5,21^.

In No-CoM trials, we found a main effect of evidence type (*X*^2^_*(1)*_ = 10.46, *p =* 0.001) showing that, on average, the P3 evoked by response-congruent evidence had bigger amplitudes than the P3 evoked by response-incongruent ones before both actions (see *Figure 2b*). In CoM trials, a significant interaction between the type of evidence and the time in the trial (*X*^2^_*(1)*_ = 7.42, *p =* 0.006) showed that the dominant amplitudes tended to reverse when participants changed their mind. At the time of the first action (*E1*), the final P3 amplitudes evoked by competing evidence numerically favoured the option selected at Action 1, although they did not differ significantly (M_Response-Congruent_ = 4.47 uV, SE = 0.47; M _Response-Incongruent =_ 4.01 uV, SE = 0.40_;_ *X*^2^_*(1)*_ = 1.22, *p =* 0.26). At the time of the second Action (*E2*), the P3 amplitude evoked by response-congruent evidence was significantly higher than that evoked by response-incongruent evidence (*X*^2^_*(1)*_ = 8.45, *p =* 0.003; see *Figure 2e;* (see *Table S1* for full statistical details).

We additionally used a simple P3-based heuristic to classify choices on each action, for each trial. In particular, we calculated the difference between the last P3 evoked by alternative evidence items (e.g. *bd* minus *pq*) and classified the trial according to the obtained sign (i.e. *bd-pq* < 0 = Pink; *bd-pq* >0 = Blue), for each condition and for CoM and No-CoM trials. We found that this simple heuristic was a significant predictor of choices (*X*^2^_*(1)*_ = 21.80, *p* <0.001), although a 4-way interaction revealed that predictions of Action 1 choices were not significant in Informative CoM trials (see *Table S2* for full statistical details). We repeated this analysis using the continuous, signed difference between competing evidence items instead of the binary heuristic. We found a significant linear relationship between the magnitude of the difference and the probability of participants’ choices (*X*^2^_*(1)*_ = 14.00, *p* <0.001; *Figure 2j*).

These results show that when a participant chose Blue at Action 1, for example, the P3 evoked by the last *bd* letter was larger than that evoked by the last *pq* (*E1*). Then, if they changed their mind and chose Pink at Action 2, the P3 evoked by the last *pq* item was larger than that evoked by the last *bd* just before Action 2 (*E2;* see *Figure 2e*).

#### 2.2.2. The P3 amplitude reflects urgency-related modulations

The first action participants made in each trial involved a level of urgency. Accumulation-to-bound theories of decision-making suggest that urgent scenarios require reducing the baseline-to-threshold distance. This can be achieved by increasing the baseline activation, lowering the threshold, increasing gain or a combination of all these mechanisms.

In No-CoM trials, the initial P3 amplitude at the start of the trial (*S1*) for urgent actions was indeed significantly higher than P3 amplitude immediately after the first action – i.e. the P3 amplitude at the starting point of the slow decision-making process leading to Action 2 (*S2*) (*X*^2^_*(1)*_ = 17.77, *p <* 0.001; see *Table S3*). Further, the final P3 amplitude reached before execution of the second action (*E2*) was smaller than that reached before the first action (*E1*) (*X*^2^_*(1)*_ = 8.15, *p =* 0.004; see *Figure 3a; Table S1*). This suggests that urgent decision-making processes started at higher baseline activation than slow, deliberate actions, and that fast decisions were achieved by increasing initial activity rather than decreasing the activity threshold.

**Figure 3.**
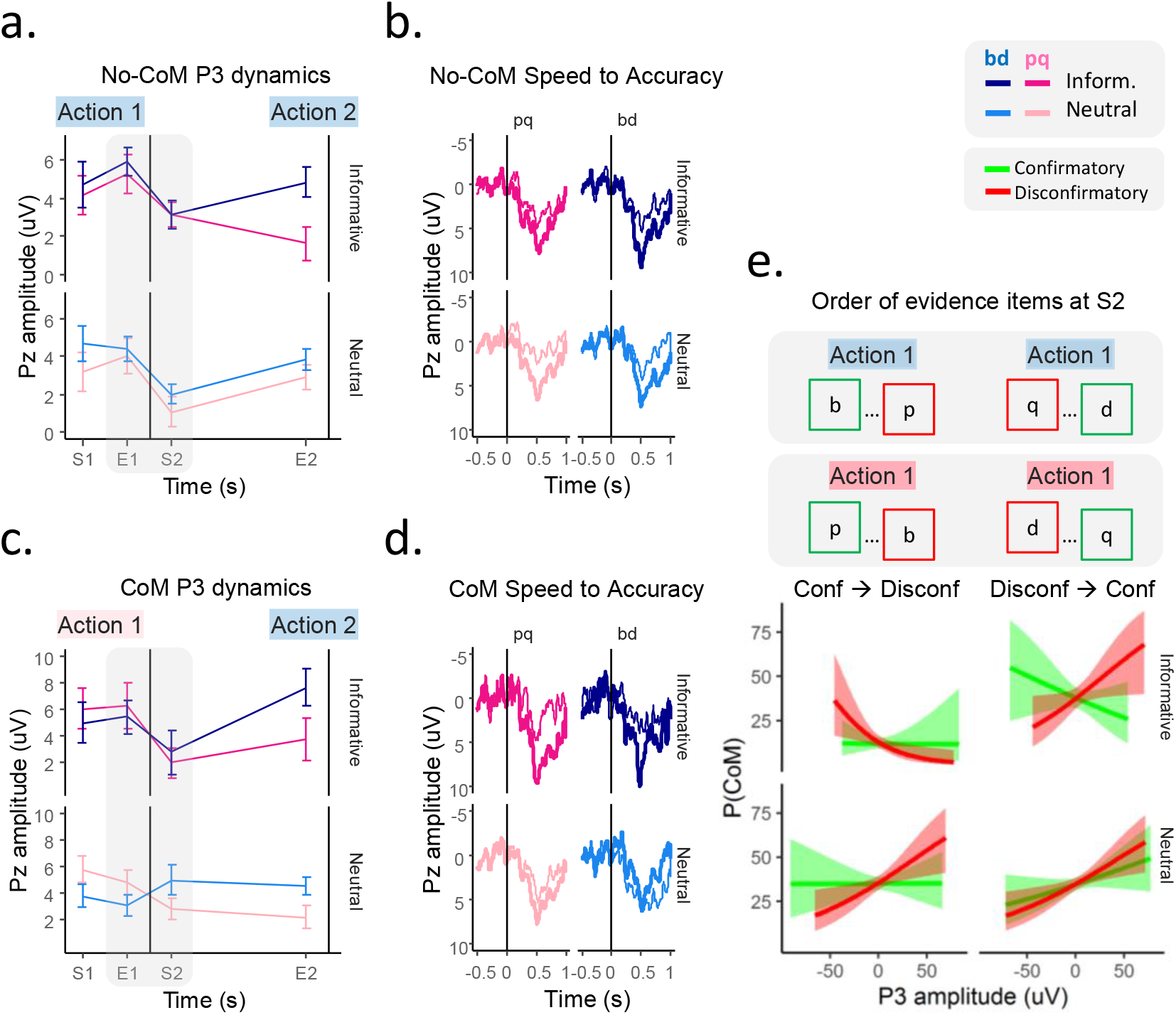
Changes in P3 amplitude track the endogenous transition from a speed to an accuracy regime. Trials in which participants chose the same colour in both Action 1 and Action 2 are illustrated on the top row (No-CoM), and those where they changed their mind in the bottom row (CoM). The graphs describe the full dataset. For illustration purposes, No-CoM trials (***a-b***) are color-coded as if participants had always chosen ‘Blue’. Similarly, CoM trials (***c-d***) are illustrated as if participants had always chosen ‘Pink’ at Action 1 and ‘Blue’ at Action 2. ***a,d***. Line graphs illustrate the grand averaged (+SEM) amplitude of the P3 evoked by the first (*S1, S2*) and last (*E1, E2*) piece of evidence presented during the decision-making processes leading to Action 1 or Action 2, for both types of evidence. ***b,d.*** Grand-averaged ERP traces at Pz, locked to the last piece of evidence presented immediately before (*E1*, *thick line*) and after (*S2*, *thin line*) Action 1. In No-CoM trials (***b***) the amplitude of the P3 significantly decreased after execution of the first action in all conditions and all types of evidence. In CoM trials (***d***), this decrease occurred in the Informative condition only. ***e.*** Probability that participants would change their mind as a function of the P3 amplitude evoked by confirmatory (*green*) and disconfirmatory (*red)* evidence items presented immediately after Action 1, sorted according to the order in which evidence items were presented. High P3 responses evoked by disconfirmatory evidence correlated with an increased probability that participants would change their mind in the Neutral condition, and in some Informative trials.

We further investigated the transition from a speed policy (Action 1) to an accuracy policy (Action 2) by comparing the P3 amplitude evoked by the last piece of evidence before the first action (*E1*) to the P3 amplitude evoked by the first evidence item presented in the accuracy regime – following Action 1 (*S2;* see *Figure 3b*). In No-Com trials, P3 amplitude drastically decreased immediately after the first action (*X*^2^_*(1)*_ = 36.21, *p <* 0.001; see *Table S4* for full statistical details). We refer to this effect as “baseline resetting”: after an initial quick decision, responses to evidence do not simply continue to accumulate. Rather, evidence processing is effectively re-set at a starting point for a second, slow evidence accumulation process.

#### 2.2.3. The P3 amplitude evoked by disconfirmatory evidence at the start of the post-decisional accumulation process predict CoM

These P3 dynamics around the time of action were altered on trials where participants changed their mind (*Figure 3c,d*). Unlike in No-CoM trials, when participants changed their mind the “baseline resetting” effect between *E1* and *S2* varied between conditions (*X*^2^_*(1)*_ = 4.76, *p =* 0.029) and evidence types (*X*^2^_*(1)*_ = 4.14, *p =* 0.041). In the Informative condition, a drop in P3 amplitude for all types of evidence again marked the transition from a speed to an accuracy regime, as in No-CoM trials. In contrast, the P3 amplitudes in Neutral conditions did not significantly decrease, overall, after Action 1 (*Figure 3c,d*). The first post-decision target letter that confirmed the preceding action had a P3 amplitude tended to decrease relative to the level immediately before the first action (*X*^2^_*(1)*_ = 2.91, *p =* 0.08). In contrast, the P3 amplitude to the first post-decision target letter disconfirming the first action did not decrease relative to *E1* (*X*^2^_*(1)*_ = 2.27, *p =* 0.13), and was in fact numerically higher than before the first action. This differential modulation resulted in a crossover effect at the transition between Action 1 and Action 2 (*Figure 3c*). These specific differences cannot be explained as a function of the P3 reaching smaller amplitudes by the time of Action 1 in CoM compared to No-CoM trials, nor to differences in the timing of actions, evidence presentation or the amount and balance of evidence seen by participants *(Supplementary Note 1)*.

We further investigated whether the P3 amplitudes evoked at the start of the post-decisional accumulation (*S2*) by confirmatory or disconfirmatory evidence specifically could predict CoM. We included the order in which confirmatory (i.e. evidence supporting the initial choice) and disconfirmatory (i.e. evidence against the initial choice) evidence items were presented after Action 1 as a control factor. An interaction between condition and P3 amplitude (*X*^2^_*(1)*_ = 6.16, *p =* 0.013) revealed that the responses evoked by disconfirmatory evidence predicted subsequent changes of mind in Neutral trials only (*X*^2^_*(1)*_ = 14.33, *p <* 0.001), regardless of the order of evidence presentation. The smaller the P3 response evoked by post-action disconfirmatory evidence, the less likely participants were to change their mind (*Figure 3e*). In Informative trials, the order of evidence presentation after an initial action significantly predicted the probability of CoM (*X*^2^_*(1)*_ = 109.17, *p <* 0.001). Participants were more likely to change their mind if the first evidence item they saw after Action 1 was disconfirmatory. In Neutral trials, the order of evidence presentation after Action 1 did not influence the CoM probability (*X*^2^_*(1)*_ = 0.04, *p =* 0.834; see *Table S5* for full statistical details).

#### 2.2.3. The P3 amplitudes at the end of trial predict confidence ratings

Finally, we aimed to test whether the P3 amplitudes contained information not only about the first order decision but also the second-order metacognitive judgement. Given that the magnitude of the P3 differences at the time of action correlated with choice probability (*Figure 2g*), we focussed on that marker (P3 difference) as a potential index of neural confidence. Specifically, we investigated whether the signed difference in P3 amplitudes evoked by the last evidence items before each action (*E1, E2)* and before letter stream stopped at the end of the trial (i.e. after Action 2) might predict subjective confidence ratings. We included end of trial amplitudes in addition to pre-action ones because post-decisional evidence can still influence confidence ratings ^22,23^, and because perceptual representations may decay over time.

Generally, participants were more confident in Informative trials than Neutral ones (*X*^2^_*(1)*_ = 65.19, *p <* 0.001), and they reported lower confidence in CoM compared to No-CoM trials (*X*^2^_*(1)*_ = 12.11, *p <* 0.001). Further, the signed difference in P3 amplitudes before Action 1 (*X*^2^_*(1)*_ = 0.70, *p =* 0.401) and Action 2 (*X*^2^_*(1)*_ = 1.23, *p =* 0.266) did not significantly predict confidence ratings. Instead, the difference evoked by the last evidence items before trial end did (*X*^2^_*(1)*_ = 9.38, *p =* 0.002). The larger the difference evoked by the last two competing evidence items before the end of the trial, the more confidence participants reported (*Figure S2a*). A follow-up analysis revealed that this effect was driven by the P3 amplitude evoked by response-incongruent evidence (*X*^2^_*(1)*_ = 9.93, *p =* 0.001): the lower the P3 amplitudes evoked by it, the higher the confidence ratings (*Figure S2b*). The P3 amplitude evoked by response-congruent evidence at the end of the trial did not significantly predict confidence ratings (*X*^2^_*(1)*_ = 0.89, *p =* 0.34; see *Table S6* for full statistical details).

## 5.3. Discussion

The capacity to revise one’s own decisions is essential to successful voluntary control of behaviour. However, understanding how varying internal states affect the processing of external evidence during decision-making and tracking its neural correlates remains a challenge. Here, we identify an EEG marker of unfolding decisional processes, and highlight the critical impact of internal urgency and confidence signals on evidence processing and their potential role in facilitating changes of mind.

### A marker of changes of mind

Previous studies have identified centro-parietal signals that reach a constant value at the decision threshold, and have a slope that correlates with the strength of evidence, thus tracking decision-making processes^18–20^. Further advancing these findings, parietal EEG signals in humans been only recently been used to make signed predictions about *which* decision participants will eventually make ^21,24^ and track how alternative choices compete during choice formation ^21^. In our task, we used sequential stimulus presentations that supported either one or another choice. Crucially, this granular approach allowed us to record the P3 evoked by each evidence item, which provided successive snapshots of the decision process as it unfolds and reflected the trajectories of each alternative. We predicted that, when participants changed their mind, the pattern of P3 amplitudes evoked by competing evidence items should reverse. That is, for example, if participants chose Pink in Action 1, the P3 evoked by response-congruent evidence (*pq)* would be greater than that to response-incongruent evidence (*bd*) in the build-up to Action 1. If they then changed their mind, to choose Blue in Action 2, this pattern should reverse, so that P3 amplitude would become higher for *bd* than for *pq* by the time of the later action. The signed difference in final amplitudes before action execution confirmed the presence of the predicted switch, and significantly predicted both action choices on a single-trial level in the neutral condition (*Figure 2d-g*). This reversal is equivalent to a change of sign in drift diffusion models of changes of mind^9^, and recalls similar findings in recordings of prefrontal neurons that track changes of mind in primates^5^. Our results thus validate the P3 as a marker of categorical choice formation and show that it can be used to track the latent state of an internal accumulator, capturing unfolding of changes of mind (see *Figure 2e*; full results in *Table S1*).

### The P3 tracks decision urgency

Our trial design involved an early, speeded decision, followed by a later, more considered decision on the same continuous stimulus stream. The effects of urgency on decision making are typically studied by comparing more urgent trials with less urgent trials ^25,26^, or as linear modulations of neural activity related to the passage of time ^11–14,27,28^. However, abrupt changes in urgency in the course of a single decision process have only rarely been studied, even though the need to speed up or slow down decision processes is a common experience in everyday life. The structure of our task allowed us to show how the internal decision variable captured by the P3 reflected sudden changes in urgency during the course of a single decision.

In No-CoM trials, we observed that the initial P3 amplitudes evoked by both evidence types at the start of the first, urgent decision were significantly higher than those at the beginning of the second, slow decision. In contrast, the final amplitude reached before Action 1 was either higher (*Figure 2a*) or comparable to (*Figure 2d*) that reached for Action 2. These results are compatible with an endogenous regulation process that reduces baseline-to-threshold distance during evidence accumulation for speeded actions^29^. Specifically, our results suggest that urgent actions may be achieved by increasing an initial activation level, corresponding to the initial P3 amplitude, rather than by reducing the neural decision threshold indexed by the final P3 amplitude. Action 1 marked the transition from speed to accuracy, and this was associated with a “baseline resetting” process in which the P3 amplitude drastically dropped immediately following Action 1 (*Figure 2b*). Interestingly, the observed results resemble the drop in frontal eye field (FEF) firing rates when macaque monkeys were exogenously cued to change from speed to accuracy regimes^30^.

### A neural turning point

In neutral CoM trials, unlike all other conditions, the P3 amplitudes following the first action did not, overall, decrease (*Figure 3d*). This overall lack of ‘resetting’ critically interacted with the nature of the evidence items. The P3 evoked by evidence *confirming* the initial choice showed a general trend to decrease, while the P3 evoked by *disconfirming* evidence remained stable or numerically increased (*Figure 2d*). This resulted in a reversal of the dominant P3 amplitudes at the start of the post-decisional evidence accumulation process, which was then maintained or amplified up to the time of Action 2 (*Figure 2e*), and eventually resulted in a change of mind. Importantly, these evidence-selective modulations could not simply reflect the urgency-related modulations described above. Urgency effects are independent of evidence, and therefore they should affect neural activity related to all available options equally^26,28^. Further, given that no differences in stimuli characteristics up to the time of the first action were observed between CoM and No-CoM trials (see *Figure S1*), it seems unlikely that variations in the processing of post-Action 1 evidence were due to external factors. Instead, our results are compatible with confidence-related modulations.

### Low confidence may facilitate changes of mind

Confidence in initial choices can influence subsequent explore/exploit decisions ^31^ and can also modulate the processing of incoming evidence ^8,16,17^. While we did not ask for any measure of subjective confidence at the time of Action 1, indirect evidence supports our interpretation on the relation between early confidence and CoM in our task. First, participants reported lower final confidence ratings on CoM trials compared to No-CoM trials (*Figure 4a, Supplementary Note 1*), and the P3 amplitudes evoked at the end of each trial significantly predicted confidence judgements (*Figure S2*). Second, changes of mind often follow low confidence choices^16^. Thus, the fact that given equal amounts of evidence people changed their minds in some trials and not in others can itself be taken as a proxy of low confidence in their initial decisions.

**Figure 4.**
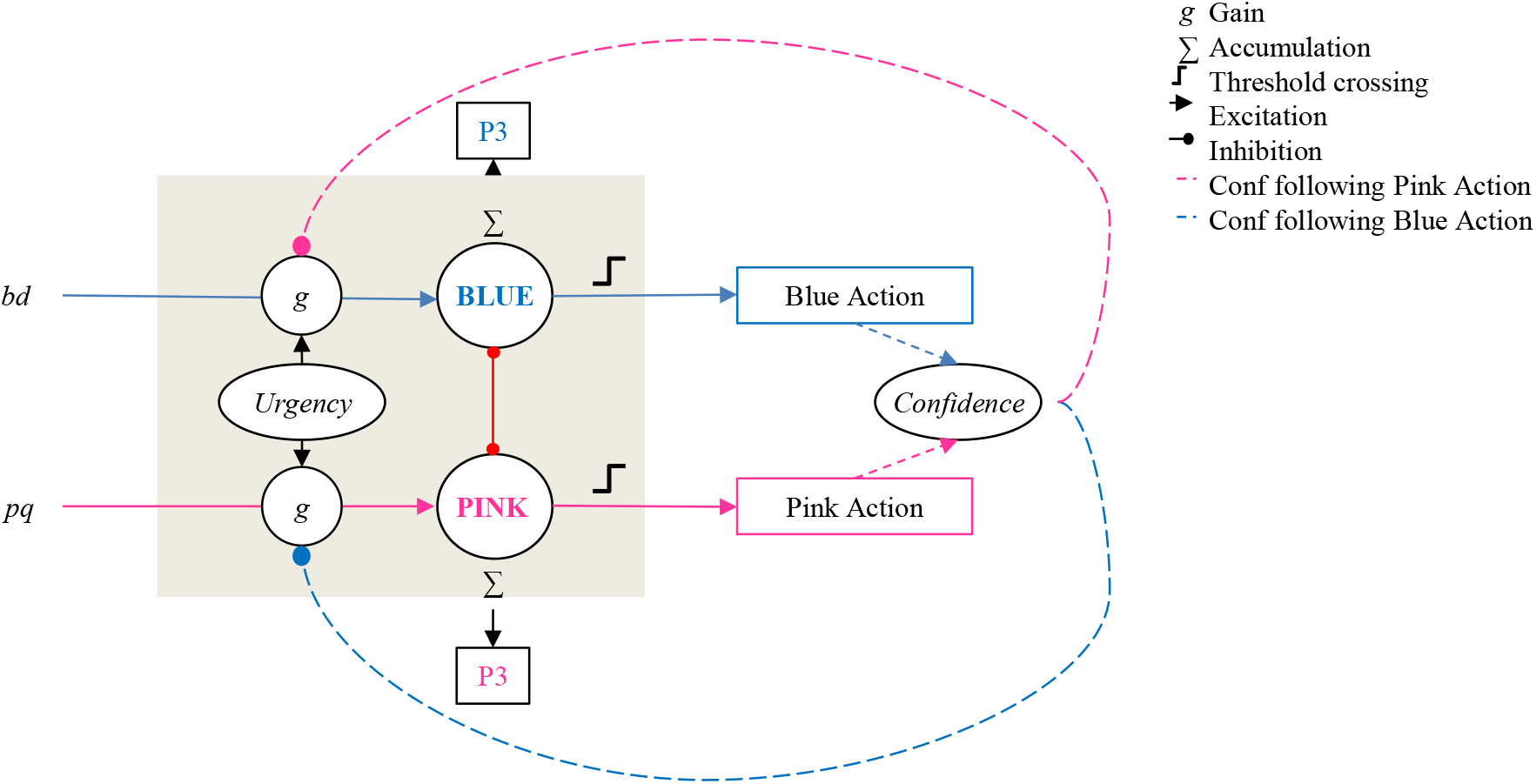
Conceptual model of the cognitive mechanisms underlying the P3. Square boxes indicate measurable events. Oval boxes indicate internal (latent) states that are not directly observable. Evidence is processed sequentially: each stimulus is processed with a certain gain (*g*), and entered into a category-specific accumulator. Based on previous research ^21,36^, we suggest that the accumulators for competing choices are mutually inhibitory. The gain in turn depends on a common urgency signal, which affects both types of evidence equally. Before the first action, urgency is high – and therefore the evoked P3 amplitudes are large from the beginning of the trial. Once one of the accumulators reaches a threshold, a decision is made. After the initial choice (Action 1), urgency decreases. This alone can account for the ‘baseline resetting’ effect observed in No-CoM trials. However, the initial quick decision in our task is assumed to be accompanied by a graded feeling of confidence. We suggest that confidence further modulates gain in the processing of subsequent evidence (*dashed arrows*). In particular, we suggest an inhibitory, asymmetric modulation. Low confidence on an initial choice (e.g. Pink action) results in an increase in gain (i.e. reduced inhibition) for the subsequent evidence disconfirming the initial choice (e.g. bd). This could account for the evidence-specific modulations of the P3 amplitude observed after Action 1 in trials where participants changed their minds. In turn, high confidence on an initial choice would result in increased inhibition of subsequent conflicting evidence accumulation.

In this study, we show a linear relationship between the neural response evoked by disconfirmatory evidence and the change of mind probability (*Figure 2j*). If the evoked P3 amplitude was small, participants were more likely to stick to their initial decision (*Figure 3f)*. Conversely, when the P3 evoked by disconfirming evidence after Action 1 was high, participants were more likely to change their mind. The P3 can thus be used as a neural marker of endogenous biases in early post-choice evidence processing. While previous research showed that a *reduction* in gain of disconfirmatory evidence makes people more prone to confirmation bias following a high-confidence choice (Rollwage et al. 2020), we here show that an *increase* in the neural processing of disconfirmatory evidence makes people more likely to change their minds, and we hypothesise that this modulation follows from low-confidence initial choices. In this sense, change of mind is the cognitive complement of confirmation bias.

Interestingly, P3s evoked by confirmatory evidence after an initial choice did not predict CoM (*Figure 3g*). Rather, neural responses to *disconfirmatory* evidence processing played the predominant role in predicting changes of mind. Thus, everything else being equal, endogenous modulations of disconfirmatory evidence processing may suffice to tilt the balance towards revision or confirmation of an initial choice. Further supporting this idea, we found that the correlation between P3 amplitudes and confidence reports was mainly driven by response-incongruent evidence at the end of the trial (*Figure S2*). Our findings contrast with previous reports that metacognitive judgements largely disregard the amount of evidence supporting un-chosen alternatives ^32–34^. However, the structure and reward system of our task readily allowed participants to change their mind and thus effectively encouraged integrating disconfirmatory evidence perhaps to a larger extent than previously used tasks.

Our finding that the P3 can reflect both first- and second-order information is in agreement both with previous findings in monkeys ^22^ and humans ^24^, and also with the idea that a single cognitive mechanism may underlie first order decisions and metacognitive judgements ^35^.

In sum, we believe that participants were less certain about their initial choices in CoM compared to No-CoM trials, and that this low confidence modulated how post-action evidence was processed. In particular, low confidence may have enhanced the neural response to disconfirmatory evidence thereby making changes of mind more likely. Figure 4 shows a conceptual model of the suggested endogenous and exogenous factors underlying the evolution of the P3 in our study.

## Conclusion

The capacity to change one’s mind is essential to successful voluntary control of behaviour. Decisions typically involve the integration of stimulus evidence and internal states, but few previous studies have been able to study the dynamics of how these two elements are integrated. Here, we identified a new neural marker that can be used as an empirical, quantifiable measure of how internal and external sources of information interact during dynamic decision-making, and we show that an endogenously-triggered boost in processing of disconfirmatory evidence following an initial choice facilitates changes of mind.

## 4. Methods

### Participants

We aimed for a final sample size of 19 participants based on a previous study ^21^. Initially, twenty-one subjects were initially invited to a single EEG session. Two participants were excluded because they did not understand the instructions. Two participants were excluded due to excessive EEG noise in the electrode of interest (Pz). Eventually, 19 participants (10 female) were included in the study (*M*_age_ = 25.3, *SD*_age_ = 3.64; range: 21-34 years).

### Procedure & experimental structure

Participants sat in a quiet room and were presented visual stimuli on a computer monitor. The instructions for the task were first displayed on the computer screen and then verbally repeated by the experimenter. Before the experiment, participants performed a practice version of the task (5 trials) to get familiar with the task. The experiment was divided into 5 blocks, with a total of 114 trials in the whole experiment. Trials were separated by a 2 s interval and participants could take a break between blocks. There were 38 Informative trials, and 76 Neutral trials. In half of the neutral trials (n = 38) evidence was presented at a slightly higher rate than on the other half (18% vs. 14% of all letters were *bd* or *pq,* respecitvely). All conditions were randomized across the experiment. The task was programmed in Python and the PsychoPy ^37^ and Pandas ^38^ toolboxes.

### Reward

The reward scheme was designed to encourage the first action (Action 1) to be fast but not completely random, and the second one (Action 2) to be as accurate as possible. Participants could win a maximum of 25 pence in each trial. To encourage the first action to be fast, they were informed that they would lose 1 penny for each second they waited before executing the first action. However, they were also encouraged to pay attention to the evidence in that first action. Incorrect actions were penalised by subtracting 12.5 p for each. Thus, in addition to the 1p/s penalisation up to the time of the first action, participants could be penalised with −12.5 p if either the first or the second action was wrong, or −25 p if both actions were wrong.

This reward scheme was only applied to informative conditions. In neutral trials, since there was no correct or incorrect choice, participants were not rewarded. If participants failed to execute two actions in any given trial, they did not obtain any reward. Participants were given averaged information about the reward obtained in each block at its end, during the breaks.

### EEG recording

EEG was recorded from 26 scalp sites (FZ, FCZ, CZ, CPZ, PZ, POZ, FC1, FC2, C1, C2, CP1, CP2, F3, F4, F7, F8, C3, C4, CP5, CP6, FC5, FC6, P3, P4, O1, O2) using active electrodes (g.LADYbird) fixed to an EEG cap (g.GAMMAcap) according to the extended international 10/20 system. EEG data were acquired using a g.GAMMAbox and g.USBamp with a sampling frequency of 256 Hz. Signal was recorded using g.Recorder (G.tec, medical engineering GmbH, Austria). All electrodes were online referenced to the right ear lobe. Vertical and horizontal electroocular activity was recorded from electrodes above and below the right eye and on the outer canthi of both eyes.

### EEG analysis

#### Preprocessing

EEG data were processed using Matlab R2014b (MathWorks), and EEGLAB version 13.5.4b ^39^. First, scalp and eye electrodes were re-referenced to the average of two mastoid electrodes. Continuous EEG and EOG data were band-pass filtered between 0.01 Hz and 30Hz using an 8th order Butterworth filter. Then, data were down-sampled to 200 Hz. Second, EEG signals were epoched in two ways. For P3analysis, EEG signal was locked from −0.5 before to 1 s after the present tion of relev nt letters (‘p’, ‘q’, ‘b’ and ‘d’). Then, an independent component analysis (ICA) was computed on the epoched data using the EEGLAB *runica* algorithm. Vertical eye movement components were visually identified and removed from the signal. Finally, artefact rejection was performed by removing all epochs with >120μV fluctuations from baseline in the preselected channels of interest (PZ).

#### Sequential P3 analysis

To study the dynamics of evidence accumulation as encoded in the P3, we extracted the P3 from electrode site Z in response to every inst nce of relev nt evidence (‘b’, ‘d’, ‘p’ and ‘q’), and we obt ined the average amplitude of the whole duration of the component [0.3 to 0.8s post stimulus], as observed in the grand-averaged data. We used these raw values for the dynamic analysis of the evolution of the P3 amplitude over time in an action-locked manner.

For average single-trial ERP analysis, we averaged the P3 component in response to response-congruent and response-incongruent separately for each single trial up to Action 1 and Action 2 respectively, and we used these values for statistical inference.

We further directly compared the amplitudes of the first and last pieces of evidence presented at the *start* (*S*) and *end* (*E*) of each evidence accumulation process (i.e. from trial onset to Action 1 (*S1, E1*) and from Action 1 to Action 2 (*S2, E2*).

To test whether P3 amplitudes had predictive power at a single trial level, we used a simple classification approach. For each single trial, and for each action, we used the following heuristic: if the last P3 (*E1/E2*) evoked by ‘bd’ stimulus was greater than that evoked by a ‘pq’ one, ‘Blue’ response was predicted. Conversely, if the 3 evoked by the last ‘pq’ stimulus before an action was greater than that evoked by the last ‘bd’ stimulus before that same action, then ‘Pink’ response was predicted. We then fitted logistic regression to predict which action participants would make, based on the P3 based prediction (Blue/Pink), the condition (Informative/Neutral) and the action (Action 1/Action 2). Further, we repeated this analysis using the continuous difference value of the P3 magnitudes rather than its sign to test whether the predictive power was linearly related to the strength of the signal.

We further tested whether the difference between the P3 amplitudes evoked by the different evidence categories before the end of the trial predicted confidence. Confidence ratings were z-scored within each participant. In particular, we ran three linear regressions to test whether confidence ratings could be predicted from 1) the signed differences between the last response-congruent and response-incongruent P3 before Action 1, Action 2 and at the end of the trial, 2) the P3 evoked by the last response-incongruent evidence item, or 3) the P3 evoked by the last response-congruent evidence item, for each condition (Informative/Neutral) and for each type of trials (No-CoM/CoM).

We fitted all multilevel linear regressions using the *lme4* package ^40^ in R. A random intercept for each subject and each trial was included to control for individual variability in all models except for the confidence and decoding ones, which included only a subject-level intercept. Full statistical details of all model outputs can be found in the *Extended data*.

## Acknowledgements

This work was supported by a joint grant from the John Templeton Foundation and the Fetzer Institute (to P.H.). The opinions expressed in this publication are those of the author(s) and do not necessarily reflect the views of the John Templeton Foundation or the Fetzer Institute.

## Author contributions

E.P-P. and P.H conceived the experiment. E.P-P and J.H. carried out the experiment. E.P-P. analysed the results. E.P-P. and P.H. wrote the manuscript, with the approval of all authors.

## Supplementary material

### *Supplementary note 1:* Additional behavioural results

#### Response bias

In neutral conditions, participants had a small but significant bias towards selecting ‘Pink’ as their final choice (*M* = 57.06% pink, *SD* = 4.00; *t*_(18)_ = 6.53, *p <* 0.001). We attribute this bias to the fact that the ‘Pink’ key was the right one in our experimental setup, and all of our participants were right-handed.

#### Time of action

There was a significant effect of condition for the time of the first (*X*^2^_(1)_ = 43.72, *p <* 0.001) and second (*X*^2^_(2)_ = 85.85, *p <* 0.001) actions. Both actions were faster in Informative (Action 1: *M =* 4.09 s*, SD =* 1.30 s; Action 2: *M =* 18.29 s*, SD =* 2.12 s) than in Neutral trials (Action 1: *M =* 4.90 s*, SD =* 1.42 s; Action 2: *M =* 20.45s*, SD =* 2.06 s).

Further, we found a significant main effect of CoM in the second action (*X*^2^_(1)_ = 10.27, *p =* 0.001), indicating that when participants changed their mind they were slower to execute the second action (CoM: *M =* 19.74 s *, SD =* 2.38 s) than when their choice was the same as in the first action (CoM: *M =* 19.00 s*, SD =* 2.27 s). The time of the first action did not differ between CoM and No-CoM trials (*X*^2^_(1)_ = 0, *p =* 0.997).

#### Perceived time of action

For simplicity, we analysed the perceived temporal distance between Action 1 and Action 2 rather than the bias for each action individually. On average the estimated bias was negative (*M =* −0.92 s, *SD =* 2.07; *t*_(18)_ = 1.93, *p* = 0.06), indicating that actions were generally perceived to be closer together in time than they actually were, though the effect failed to reach significance.

#### Evidence at time of action

We calculated the total amount of evidence items, the balance between competing evidence items preceding each action and the average time of presentation of competing evidence types in CoM and No-CoM trials (*Figure S1*). The total amount of evidence items viewed by participants up to the time of Action 1 did not differ between conditions nor Com/No-CoM trials. We found that there were no differences between Neutral CoM and No-CoM trials in the net balance of evidence before the first Action 1 (*X*^2^_*(1)*_ = 0.23, *p =* 0.63). Instead, in Informative trials, the first action was supported by less net evidence on trials where participants changed their mind than on those where they did not (*X*^2^_*(1)*_ = 43.63, *p <* 0.001). The average time at which evidence of different kinds was presented did not vary between CoM/No-CoM trials before Action 1 (*X*^2^_*(1)*_ = 0.7759, *p =* 0.378). Further, we also found that participants viewed more evidence items before committing to a final decision (Action 2) in CoM compared to No-CoM trials (*X*^2^_*(1)*_ = 10.00, *p =* 0.001) and that at the moment of action the total balance of evidence supported the final choice more strongly (*X*^2^_*(1)*_ = 257.17, *p <* 0.001). These results suggest that change of mind was associated with a compensatory shift from a first action based on a relatively low accuracy criterion, to a second action with a more conservative criterion.

---

**Figure S1.**
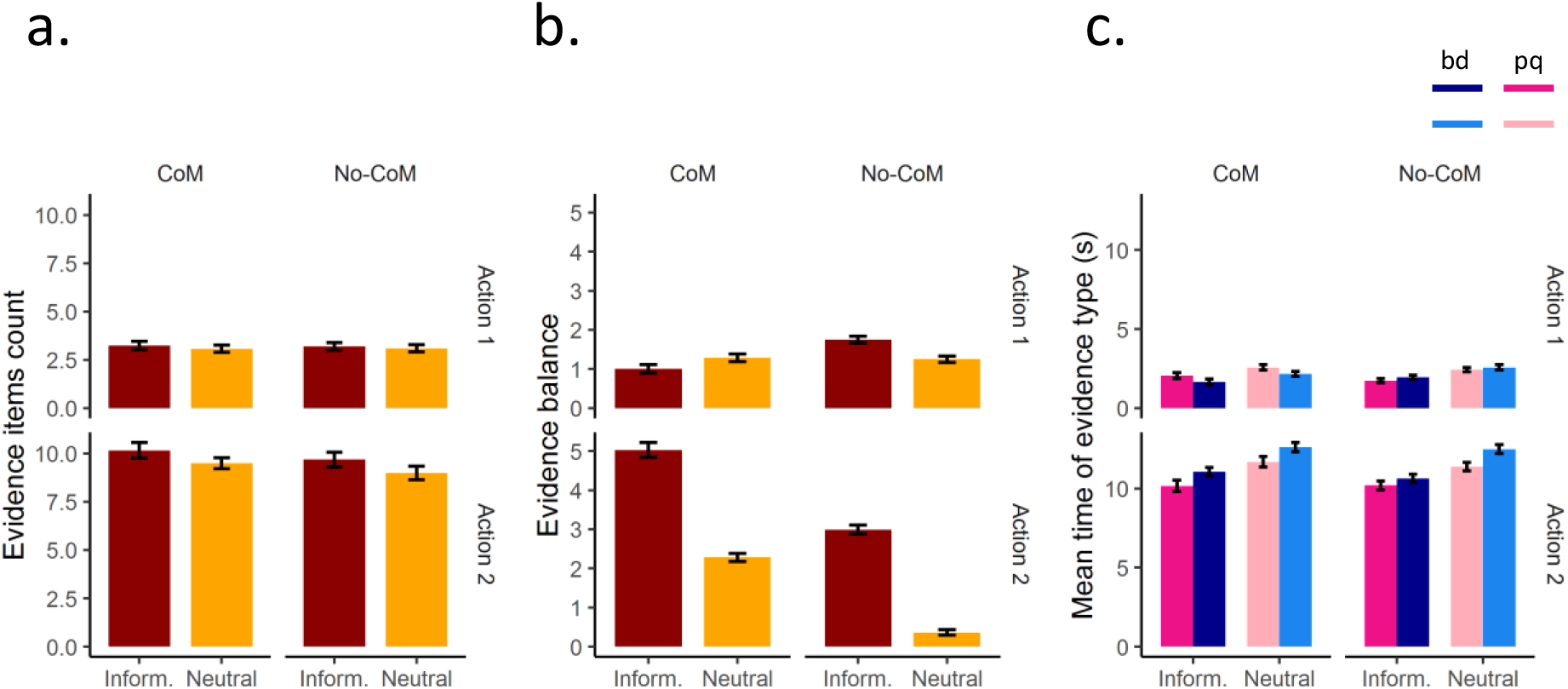
Stimulus features at the time of decision. ***a.*** Grand-average (± SEM) total amount of evidence items before Action 1 and 2 in CoM and No-CoM trials. There was no difference in the total number of evidence items viewed before the first guess (Action) 1 in CoM vs. No-CoM trials, but participants waited for more evidence before committing to a final response (Action 2) in CoM trials. ***b.*** Balance of evidence before action (response-congruent minus response-incongruent with respect to Action 1 and 2). In the Informative conditions only, participants’ first ction was supported by more net evidence in CoM trials than No-CoM ones. In CoM trials, participants committed to a final response (Action 2) when net evidence was on average much higher than that in No-CoM trials. **c.** Average time of presentation of different types of evidence before each action. Participants tended to respond shortly after an evidence item supporting their choice. This is reflected in the higher average presentation times for response-congruent evidence. No other significant differences between CoM and No-CoM trials were observed. For illustration purposes only, colour coding of CoM (Action 1: Pink, Action 2: Blue) and No-CoM (Action 1: Blue, Action 2: Blue) trials is the same as in the main manuscript.

**Figure S2.**
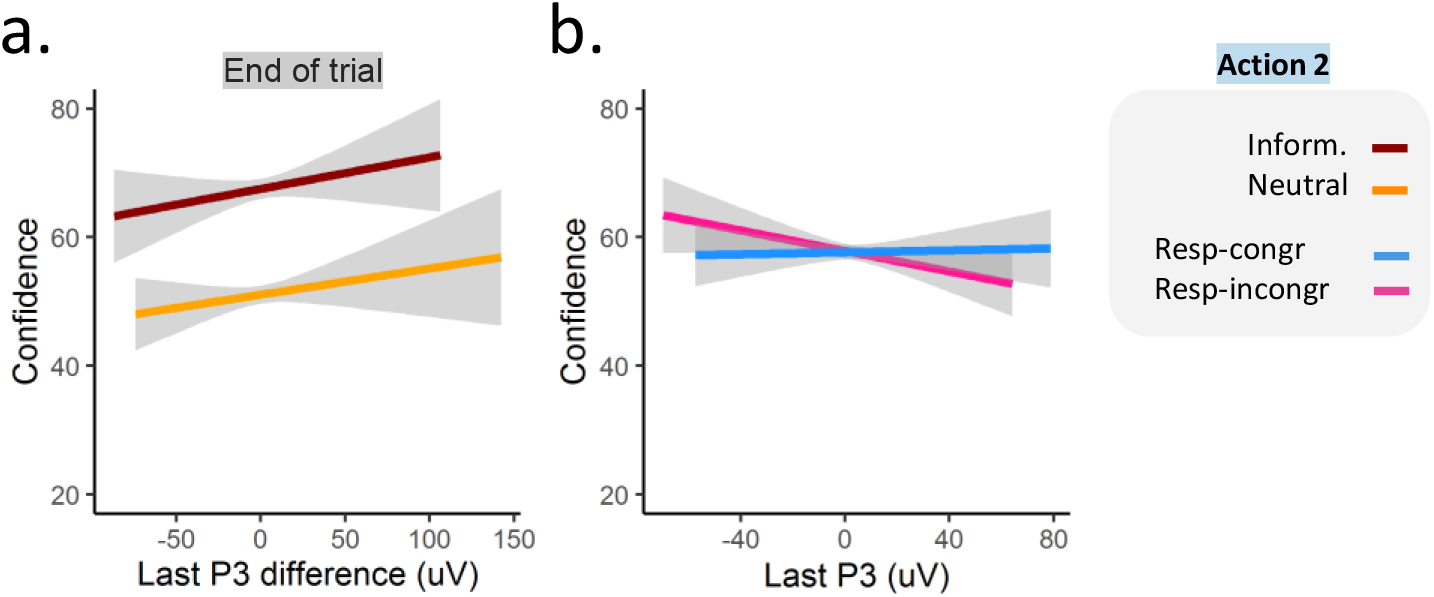
The P3 correlates with confidence ratings. The illustrated data correspond to all types trials (No-CoM & CoM). For illustration purposes, data are color-coded as if participants had chosen ‘Blue’ at Action 2 in all trials. ***a.*** The signed difference between the last P3 amplitudes (Pz electrode) evoked by final response-congruent and response-incongruent evidence (i.e. blue minus pink in this example) at the end of the trial (*right*) is plotted in relation to the confidence ratings. The greater the difference in the evoked P3s at the end of the trial, the more confidence participants reported. ***b.*** The amplitude of the P3 evoked by the last piece of response-congruent (*blue*) evidence before trial end did not predict confidence ratings. Instead, the amplitude of the P3 evoked by response-incongruent (*pink*) did. The lower the P3 amplitude evoked by the last item of response-incongruent evidence, the more confidence participants reported. Informative and Neutral conditions are pooled together.

## 5. Extended data

### 1.1. Sequential P3 analysis

**Table S1.** P3 amplitudes correlate with choice.

| A. No-CoM trials:<br>P3 ~ TimePoint*Condition *EvidenceType + (1 \| Sub/Trial);<br>TimePoint: E1/E2 |  |  |  |
| --- | --- | --- | --- |
| Fixed Factor | Chisq | Df | Pr(>Chisq) |
| **TimePoint** | **8.154222** | **1** | **0.004296** |
| Condition | 1.175409 | 1 | 0.278293 |
| **EvidenceType** | **10.46013** | **1** | **0.00122** |
| TimePoint:Condition | 3.058416 | 1 | 0.08032 |
| TimePoint:EvidenceType | 1.246411 | 1 | 0.264239 |
| Condition:EvidenceType | 2.768101 | 1 | 0.09616 |
| TimePoint:EvidenceType:Condition | 1.73899 | 1 | 0.187267 |
| B. CoM trials:<br>P3 ~ TimePoint*Condition *EvidenceType + (1 \| Sub/Trial);<br>TimePoint: E1/E2 |  |  |  |
| Fixed Factor | Chisq | Df | Pr(>Chisq) |
| TimePoint | 0.994943 | 1 | 0.318537 |
| **Condition** | **7.934492** | **1** | **0.00485** |
| EvidenceType | 1.876278 | 1 | 0.170758 |
| TimePoint:Condition | 0.023457 | 1 | 0.878274 |
| **TimePoint:EvidenceType** | **7.423443** | **1** | **0.006438** |
| Condition:EvidenceType | 0.866105 | 1 | 0.352036 |
| TimePoint:EvidenceType:Condition | 0.047487 | 1 | 0.827496 |

**Table S2.** Decoding choice.

| **A. Full model:**<br>**P3 ~ ClasOut\*Action\*Condition\*CoM + (1\|Sub);** |  |  |  |
| --- | --- | --- | --- |
| Fixed Factor | Chisq | Df | Pr(>Chisq) |
| **ClasOut** | **21.80564** | **1** | **<0.001** |
| **Action** | **10.07733** | **1** | **0.001501** |
| **Condition** | **9.788537** | **1** | **0.001756** |
| **CoM** | **7.176786** | **1** | **0.007385** |
| ClasOut:Action | 0.136785 | 1 | 0.711498 |
| ClasOut:Condition | 2.624997 | 1 | 0.105193 |
| Action:Condition | 2.447005 | 1 | 0.117749 |
| ClasOut:CoM | 0.625717 | 1 | 0.428931 |
| **Action:CoM** | **15.12794** | **1** | **0.0001** |
| Condition:CoM | 0.364567 | 1 | 0.545981 |
| ClasOut:Action:Condition | 3.322342 | 1 | 0.068344 |
| ClasOut:Action:CoM | 0.092469 | 1 | 0.761062 |
| **ClasOut:Condition:CoM** | **4.827963** | **1** | **0.028002** |
| **Action:Condition:CoM** | **10.34457** | **1** | **0.001299** |
| **ClasOut:Action:Condition:CoM** | **4.074014** | **1** | **0.043548** |
| **B. CoM trials:**<br>**P3 ~ ClasOut\*Action\*Condition + (1\|Sub);** |  |  |  |
| Fixed Factor | Chisq | Df | Pr(>Chisq) |
| ClasOut | 3.159125 | 1 | 0.075504 |
| **Action** | **26.6739** | **1** | **<0.001** |
| Condition | 1.021572 | 1 | 0.312146 |
| ClasOut:Action | 0.08842 | 1 | 0.766196 |
| ClasOut:Condition | 1.154329 | 1 | 0.282645 |
| **Action:Condition** | **12.28637** | **1** | **0.000456** |
| **ClasOut:Action:Condition** | **7.208952** | **1** | **0.007254** |
| **C. CoM trials, Action 1:**<br>**P3 ~ ClasOut\*Condition + (1\|Sub);** |  |  |  |
| Fixed Factor | Chisq | Df | Pr(>Chisq) |
| ClasOut | 0.733334 | 1 | 0.391805 |
| **Condition** | **4.412347** | **1** | **0.03568** |
| **ClasOut:Condition** | **7.648696** | **1** | **0.00568** |
| **D. CoM trials, Action 1, Informative:**<br>**P3 ~ ClasOut + (1\|Sub);** |  |  |  |
| Fixed Factor | Chisq | Df | Pr(>Chisq) |
| ClasOut | 3.841252 | 1 | 0.050006 |
| **E. CoM trials, Action 1, Neutral:**<br>**P3 ~ ClasOut + (1\|Sub);** |  |  |  |

| Fixed Factor | Chisq | Df | Pr(>Chisq) |
| --- | --- | --- | --- |
| **ClasOut** | **4.595824** | **1** | **0.03205** |

**F. No-CoM trials: P3 ~ ClasOut\*Action\*Condition + (1|Sub);**
| Fixed Factor | Chisq | Df | Pr(>Chisq) |
| --- | --- | --- | --- |
| **ClasOut** | **20.18527** | **1** | **<0.001** |
| Action | 0.148214 | 1 | 0.700248 |
| **Condition** | **9.640623** | **1** | **0.001903** |
| ClasOut:Action | 0.0479 | 1 | 0.826759 |
| **ClasOut:Condition** | **6.470966** | **1** | **0.010965** |
| Action:Condition | 0.101344 | 1 | 0.750223 |
| ClasOut:Action:Condition | 0.240756 | 1 | 0.62366 |

**G. No-CoM trials, Neutral: P3 ~ ClasOut\*Action + (1|Sub);**
| Fixed Factor | Chisq | Df | Pr(>Chisq) |
| --- | --- | --- | --- |
| **ClasOut** | **5.010671** | **1** | **0.025192** |
| Action | 0.041331 | 1 | 0.8389 |
| ClasOut:Action | 0.01404 | 1 | 0.905681 |

**H. No-CoM trials, Informative: P3 ~ ClasOut\*Action + (1|Sub);**
| Fixed Factor | Chisq | Df | Pr(>Chisq) |
| --- | --- | --- | --- |
| **ClasOut** | **22.17924** | **1** | **<0.001** |
| Action | 0.195818 | 1 | 0.658118 |
| ClasOut:Action | 0.292937 | 1 | 0.588345 |

**Table S3.** P3 amplitude at the start of evidence accumulation for Action 1 and Action 2.

| **A. Full model:**<br>**P3 ~ TimePoint\*Condition\*CoM\*EvidenceType + (1\|Sub/Trial)**<br>TimePoint: S1/S2 |  |  |  |
| --- | --- | --- | --- |
| Fixed Factor | Chisq | Df | Pr(>Chisq) |
| TimePoint | **24.80857** | **1** | **<0.001** |
| Condition | **4.95841** | **1** | **0.025964** |
| CoM | **8.735527** | **1** | **0.003121** |
| EvidenceType | 2.79278 | 1 | 0.09469 |
| TimePoint:Condition | 0.006925 | 1 | 0.93368 |
| TimePoint:CoM | 0.327359 | 1 | 0.567218 |
| Condition:CoM | 1.045092 | 1 | 0.30664 |
| TimePoint:EvidenceType | 0.447946 | 1 | 0.503312 |
| Condition:EvidenceType | 0.744086 | 1 | 0.388355 |
| CoM:EvidenceType | 1.428869 | 1 | 0.231949 |
| TimePoint:Condition:CoM | 2.85776 | 1 | 0.090934 |
| TimePoint:Condition:EvidenceType | 0.095929 | 1 | 0.756771 |
| TimePoint:CoM:EvidenceType | **5.773835** | **1** | **0.016266** |
| Condition:CoM:EvidenceType | 0.55655 | 1 | 0.455654 |
| TimePoint:Condition:CoM:EvidenceType | 0.659676 | 1 | 0.416674 |
| **B. No-CoM:**<br>**P3 ~ TimePoint\*Condition\*EvidenceType + (1\|Sub/Trial)**<br>TimePoint: S1/S2 |  |  |  |
| Fixed Factor | Chisq | Df | Pr(>Chisq) |
| TimePoint | **17.7749** | **1** | **<0.001** |
| Condition | **5.517292** | **1** | **0.018829** |
| EvidenceType | **3.900675** | **1** | **0.048267** |
| TimePoint:Condition | 0.829431 | 1 | 0.362437 |
| TimePoint:EvidenceType | 0.736439 | 1 | 0.390804 |
| Condition:EvidenceType | 1.13346 | 1 | 0.287038 |
| TimePoint:Condition:EvidenceType | 0.023643 | 1 | 0.877797 |
| **C. CoM:**<br>**P3 ~ TimePoint\*Condition\*EvidenceType + (1\|Sub/Trial)**<br>TimePoint: S1/S2 |  |  |  |
| Fixed Factor | Chisq | Df | Pr(>Chisq) |
| TimePoint | **7.292737** | **1** | **0.006923** |
| Condition | 0.042284 | 1 | 0.837079 |
| EvidenceType | 0.005196 | 1 | 0.942534 |
| TimePoint:Condition | 2.083071 | 1 | 0.14894 |
| TimePoint:EvidenceType | **6.571816** | **1** | **0.010361** |
| Condition:EvidenceType | 0.02828 | 1 | 0.866453 |
| TimePoint:Condition:EvidenceType | 0.824749 | 1 | 0.363795 |

**Table S4.** Speed to accuracy transition.

| **A. Full model:**<br>**P3 ~ TimePoint\*EvidenceType\*CoM\*Condition + (1\|Sub/Trial)**<br>TimePoint: E1/S2 |  |  |  |
| --- | --- | --- | --- |
| Fixed Factor | Chisq | Df | Pr(>Chisq) |
| **TimePoint** | **36.23502** | **1** | **<0.001** |
| EvidenceType | 2.226966 | 1 | 0.13562 |
| **CoM** | **4.037636** | **1** | **0.044496** |
| **Condition** | **9.976427** | **1** | **0.001586** |
| TimePoint:EvidenceType | 0.965845 | 1 | 0.325718 |
| **TimePoint:CoM** | **4.13057** | **1** | **0.042115** |
| EvidenceType:CoM | 0.233495 | 1 | 0.628945 |
| TimePoint:Condition | 1.241225 | 1 | 0.265235 |
| EvidenceType:Condition | 0.272066 | 1 | 0.601949 |
| CoM:Condition | 0.866667 | 1 | 0.35188 |
| TimePoint:EvidenceType:CoM | 2.693017 | 1 | 0.100789 |
| TimePoint:EvidenceType:Condition | 0.646772 | 1 | 0.421269 |
| TimePoint:CoM:Condition | 3.612441 | 1 | 0.057349 |
| EvidenceType:CoM:Condition | 0.252409 | 1 | 0.615384 |
| TimePoint:EvidenceType:CoM:Condition | 0.24839 | 1 | 0.618211 |
| **B. No-CoM trials:**<br>**P3 ~ TimePoint\*EvidenceType\*Condition + (1\|Sub/Trial)**<br>TimePoint: E1/S2 |  |  |  |
| Fixed Factor | Chisq | Df | Pr(>Chisq) |
| **TimePoint** | **36.21953** | **1** | **<0.001** |
| EvidenceType | 2.206744 | 1 | 0.137408 |
| **Condition** | **10.09537** | **1** | **0.001486** |
| TimePoint:EvidenceType | 0.051588 | 1 | 0.820322 |
| TimePoint:Condition | 5.05E-05 | 1 | 0.994331 |
| EvidenceType:Condition | 0.480721 | 1 | 0.488096 |
| TimePoint:EvidenceType:Condition | 0.175851 | 1 | 0.674964 |
| **C. CoM trials:**<br>**P3 ~ TimePoint\*EvidenceType\*Condition + (1\|Sub/Trial)**<br>TimePoint: E1/S2 |  |  |  |
| Fixed Factor | Chisq | Df | Pr(>Chisq) |
| TimePoint | 3.182456 | 1 | 0.074433 |
| EvidenceType | 0.271609 | 1 | 0.602254 |
| Condition | 0.537653 | 1 | 0.463407 |
| **TimePoint:EvidenceType** | **4.146313** | **1** | **0.041725** |
| **TimePoint:Condition** | **4.766559** | **1** | **0.029018** |
| EvidenceType:Condition | 0.029033 | 1 | 0.864703 |
| TimePoint:EvidenceType:Condition | 0.760223 | 1 | 0.383259 |

**Table S5.** Evidence processing after Action 1.

| **A. Full model:**<br>**P(CoM) ~ P3\*EvidenceType\*Condition\*EvidenceOrder + (1\|Sub)** |  |  |  |
| --- | --- | --- | --- |
| Fixed Factor | Chisq | Df | Pr(>Chisq) |
| **P3** | **8.925681** | **1** | **0.002812** |
| EvidenceType | 0.436445 | 1 | 0.508843 |
| **Condition** | **42.26754** | **1** | **<0.001** |
| **EvidenceOrder** | **27.86017** | **1** | **<0.001** |
| **P3:EvidenceType** | **4.08141** | **1** | **0.043357** |
| **P3:Condition** | **5.75647** | **1** | **0.016428** |
| EvidenceType:Condition | 0.072297 | 1 | 0.788021 |
| P3:EvidenceOrder | 2.448071 | 1 | 0.117669 |
| EvidenceType:EvidenceOrder | 0.014571 | 1 | 0.903921 |
| **Condition:EvidenceOrder** | **74.60875** | **1** | **<0.001** |
| P3:EvidenceType:Condition | 0.159873 | 1 | 0.689273 |
| P3:EvidenceType:EvidenceOrder | 1.298896 | 1 | 0.254415 |
| P3:Condition:EvidenceOrder | 1.610032 | 1 | 0.204487 |
| EvidenceType:Condition:EvidenceOrder | 0.018503 | 1 | 0.8918 |
| **P3:EvidenceType:Condition:EvidenceOrder** | **7.988059** | **1** | **0.004709** |
| **B. Neutral trials:**<br>**P(CoM) ~ P3\*EvidenceType\*EvidenceOrder + (1\|Sub)** |  |  |  |
| Fixed Factor | Chisq | Df | Pr(>Chisq) |
| **P3** | **14.0988** | **1** | **0.000173** |
| EvidenceType | 0.188924 | 1 | 0.663814 |
| EvidenceOrder | 0.029636 | 1 | 0.86332 |
| **P3:EvidenceType** | **4.072936** | **1** | **0.043575** |
| P3:EvidenceOrder | 0.674171 | 1 | 0.411601 |
| EvidenceType:EvidenceOrder | 0.022581 | 1 | 0.880551 |
| P3:EvidenceType:EvidenceOrder | 0.02452 | 1 | 0.875568 |
| **C. Neutral trials, confirmatory evidence:**<br>**P(CoM) ~ P3\*EvidenceOrder + (1\|Sub)** |  |  |  |
| Fixed Factor | Chisq | Df | Pr(>Chisq) |
| P3 | 1.293897 | 1 | 0.255331 |
| EvidenceOrder | 0.047414 | 1 | 0.827625 |
| P3:EvidenceOrder | 0.552963 | 1 | 0.45711 |
| **D. Neutral trials, disconfirmatory evidence:**<br>**P(CoM) ~ P3\*EvidenceOrder + (1\|Sub)** |  |  |  |
| Fixed Factor | Chisq | Df | Pr(>Chisq) |
| **P3** | **16.89406** | **1** | **<0.001** |
| EvidenceOrder | 0.002817 | 1 | 0.957673 |
| P3:EvidenceOrder | 0.170875 | 1 | 0.679335 |

| E. Informative trials:<br>P(CoM) ~ P3*EvidenceType*EvidenceOrder + (1\|Sub) |  |  |  |
| --- | --- | --- | --- |
| Fixed Factor | Chisq | Df | Pr(>Chisq) |
| P3 | 0.139665 | 1 | 0.708614 |
| EvidenceType | 0.33016 | 1 | 0.565565 |
| **EvidenceOrder** | **106.0005** | **1** | **<0.001** |
| P3:EvidenceType | 0.291757 | 1 | 0.589097 |
| P3:EvidenceOrder | 3.60002 | 1 | 0.057779 |
| EvidenceType:EvidenceOrder | 0.004509 | 1 | 0.946464 |
| **P3:EvidenceType:EvidenceOrder** | **8.521094** | **1** | **0.003511** |
| F. Informative trials, confirmatory evidence:<br>P(CoM) ~ P3*EvidenceOrder + (1\|Sub) |  |  |  |
| Fixed Factor | Chisq | Df | Pr(>Chisq) |
| P3 | 0.625578 | 1 | 0.428982 |
| **EvidenceOrder** | **54.36083** | **1** | **<0.001** |
| P3:EvidenceOrder | 0.444819 | 1 | 0.504806 |
| G. Informative trials, disconfirmatory evidence:<br>P(CoM) ~ P3*EvidenceOrder + (1\|Sub) |  |  |  |
| Fixed Factor | Chisq | Df | Pr(>Chisq) |
| P3 | 0.040129 | 1 | 0.841229 |
| **EvidenceOrder** | **50.43971** | **1** | **<0.001** |
| **P3:EvidenceOrder** | **11.42279** | **1** | **0.000725** |
| H. Informative trials, disconfirmatory evidence, disconfirmatory evidence first:<br>P(CoM) ~ P3 + (1\|Sub) |  |  |  |
| Fixed Factor | Chisq | Df | Pr(>Chisq) |
| **P3** | **7.081036** | **1** | **0.00779** |
| I. Informative trials, disconfirmatory evidence, confirmatory evidence first:<br>P(CoM) ~ P3 + (1\|Sub) |  |  |  |
| Fixed Factor | Chisq | Df | Pr(>Chisq) |
| **P3** | **4.196063** | **1** | **0.040518** |

**Table S6.** P3-based confidence models.

| **A. P3 signed difference before Action 1:**<br>**zConfidence ~ P3diff\*CoM\*Condition + (1\|Sub)** |  |  |  |
| --- | --- | --- | --- |
| Fixed Factor | Chisq | Df | Pr(>Chisq) |
| P3diff | 0.703955 | 1 | 0.401458 |
| **Condition** | **241.5759** | **1** | **<0.001** |
| **CoM** | **26.02095** | **1** | **<0.001** |
| P3diff:Condition | 0.577273 | 1 | 0.447383 |
| P3diff:CoM | 0.013405 | 1 | 0.907828 |
| Condition:Congruenece | 1.139452 | 1 | 0.285768 |
| P3diff:Condition:CoM | 0.074904 | 1 | 0.784326 |
| **B. P3 signed difference before Action 2:**<br>**zConfidence ~ P3diff\*CoM\*Condition + (1\|Sub)** |  |  |  |
| Fixed Factor | Chisq | Df | Pr(>Chisq) |
| P3diff | 1.233327 | 1 | 0.266761 |
| **Condition** | **408.9921** | **1** | **<0.001** |
| **CoM** | **34.6443** | **1** | **<0.001** |
| P3diff:Condition | 1.544844 | 1 | 0.213898 |
| P3diff:CoM | 0.054144 | 1 | 0.816004 |
| Condition:Congruenece | 2.566329 | 1 | 0.109161 |
| P3diff:Condition:CoM | 0.512151 | 1 | 0.474209 |
| **C. P3 signed difference before trial end:**<br>**zConfidence ~ P3diff\*CoM\*Condition + (1\|Sub)** |  |  |  |
| Fixed Factor | Chisq | Df | Pr(>Chisq) |
| **P3diff** | **9.386098** | **1** | **0.002186** |
| **Condition** | **286.3893** | **1** | **<0.001** |
| **CoM** | **24.86162** | **1** | **<0.001** |
| P3diff:Condition | 1.712217 | 1 | 0.190698 |
| P3diff:CoM | 1.750255 | 1 | 0.185845 |
| Condition:Congruenece | 0.178804 | 1 | 0.672403 |
| P3diff:Condition:CoM | 0.059194 | 1 | 0.807774 |
| **D. Last response-congruent P3 before trial end:**<br>**zConfidence ~ P3_RCongr\*CoM\*Condition + (1\|Sub)** |  |  |  |
| Fixed Factor | Chisq | Df | Pr(>Chisq) |
| P3_RCongr | 0.92122 | 1 | 0.337155 |
| **Condition** | **284.369** | **1** | **<0.001** |
| **CoM** | **25.059** | **1** | **<0.001** |
| P3_RCongr:Condition | 0.177459 | 1 | 0.673566 |
| P3_RCongr:CoM | 0.01921 | 1 | 0.889767 |
| Condition:Congruenece | 0.179881 | 1 | 0.671476 |
| P3_RCongr:Condition:CoM | 2.276296 | 1 | 0.131365 |

| E. Last response-incongruent P3 before trial end:<br>zConfidence ~ P3_RIncongr*CoM*Condition + (1\|Sub) |  |  |  |
| --- | --- | --- | --- |
| Fixed Factor | Chisq | Df | Pr(>Chisq) |
| **P3_RIncongr** | **9.884232** | **1** | **0.001667** |
| **Condition** | **286.3055** | **1** | **<0.001** |
| **CoM** | **25.26583** | **1** | **<0.001** |
| P3_RIncongr:Condition | 2.209769 | 1 | 0.137139 |
| P3_RIncongr:CoM | 2.926146 | 1 | 0.087155 |
| Condition:Congruenece | 0.159394 | 1 | 0.689715 |
| P3_RIncongr:Condition:CoM | 3.604904 | 1 | 0.057609 |

## References

1. Nickerson, R. S. Confirmation bias: a ubiquitous phenomenon in many guises. Rev. Gen. Psychol. 2, 175–220 (1998).

2. Forstmann, B. U., Ratcliff, R. & Wagenmakers, E. Sequential Sampling Models in Cognitive Neuroscience: Advantages, Applications, and Extensions. Annu. Rev. Psychol. 67, 641–666 (2016).

3. Ratcliff, R., Smith, P. L., Brown, S. D. & McKoon, G. Diffusion Decision Model: Current Issues and History. Trends Cogn. Sci. 20, 260–281 (2016).

4. Resulaj, A., Kiani, R., Wolpert, D. M. & Shadlen, M. N. Changes of mind in decision-making. Nature 461, 263–6 (2009).

5. Kiani, R., Cueva, C. J., Reppas, J. B. & Newsome, W. T. Dynamics of neural population responses in prefrontal cortex indicate changes of mind on single trials. Curr. Biol. (2014) doi:10.1016/j.cub.2014.05.049.

6. Albantakis, L., Branzi, F. M., Costa, A. & Deco, G. A multiple-choice task with changes of mind. PLoS One 7, (2012).

7. O’Connell, R. G. & Murphy, P. R. U-turns in the brain. Nat. Neurosci. 21, 461–462 (2018).

8. Fleming, S. M., Van Der Putten, E. J. & Daw, N. D. Neural mediators of changes of mind about perceptual decisions. Nat. Neurosci. 21, 617–624 (2018).

9. Resulaj, A., Kiani, R., Wolpert, D. M. & Shadlen, M. N. Changes of mind in decision-making. Nature 461, 263–6 (2009).

10. Wickelgren, W. A. Speed-accuracy tradeoff and information processing dynamics. Acta Psychol. (Amst). 41, 67–85 (1977).

11. Thura, D. & Cisek, P. Modulation of Premotor and Primary Motor Cortical Activity during Volitional Adjustments of Speed-Accuracy Trade-Offs. J. Neurosci. 36, 938–956 (2016).

12. Thura, D., Cos, I., Trung, J. & Cisek, P. Context-dependent urgency influences speed-accuracy trade-offs in decision-making and movement execution. J. Neurosci. 34, 16442–16454 (2014).

13. Cisek, P., Puskas, G. a. & El-Murr, S. Decisions in Changing Conditions: The Urgency-Gating Model. J. Neurosci. 29, 11560–11571 (2009).

14. Thura, D. & Cisek, P. Deliberation and commitment in the premotor and primary motor cortex during dynamic decision making. Neuron 81, 1401–16 (2014).

15. Rollwage, M., Dolan, R. J. & Fleming, S. M. Metacognitive Failure as a Feature of Those Holding Radical Beliefs. Curr. Biol. 28, 4014–4021.e8 (2018).

16. Rollwage, M. et al. Confidence drives a neural confirmation bias. Nat. Commun. 11, (2020).

17. Balsdon, T., Wyart, V. & Mamassian, P. Confidence controls perceptual evidence accumulation. Nat. Commun. 11, (2020).

18. Twomey, D. M., Murphy, P. R., Kelly, S. P. & O’Connell, R. G. The classic P300 encodes a build-to-threshold decision variable. Eur. J. Neurosci. 42, 1636–1643 (2015).

19. O’Connell, R. G., Dockree, P. M. & Kelly, S. P. A supramodal accumulation-to-bound signal that determines perceptual decisions in humans. Nat. Neurosci. (2012) doi:10.1038/nn.3248.

20. Kelly, S. P. & O’Connell, R. G. Internal and External Influences on the Rate of Sensory Evidence Accumulation in the Human Brain. J. Neurosci. 33, 19434–19441 (2013).

21. Parés-Pujolràs, E., Travers, E., Ahmetoglu, Y. & Haggard, P. Evidence accumulation under uncertainty – a neural marker of emerging choice and urgency. 1–49 (2020).

22. Kiani, R. & Shadlen, M. N. Representation of confidence associated with a decision by neurons in the parietal cortex. Science 324, 759–64 (2009).

23. Kiani, R., Corthell, L. & Shadlen, M. N. Choice certainty is informed by both evidence and decision time. Neuron (2014) doi:10.1016/j.neuron.2014.12.015.

24. Herding, J., Ludwig, S., von Lautz, A., Spitzer, B. & Blankenburg, F. Centro-parietal EEG potentials index subjective evidence and confidence during perceptual decision making. Neuroimage 201, 116011 (2019).

25. Reddi, B. A. J. & Carpenter, R. H. S. The influence of urgency on decision time. Nat. Neurosci. 3, 827–830 (2000).

26. Steinemann, N. A., O’Connell, R. G. & Kelly, S. P. Decisions are expedited through multiple neural adjustments spanning the sensorimotor hierarchy. Nat. Commun. 9, 3627 (2018).

27. Thura, D. Decision urgency invigorates movement in humans. Behav. Brain Res. 382, 112477 (2020).

28. Churchland, A. K., Kiani, R. & Shadlen, M. N. Decision-making with multiple alternatives. Nat. Neurosci. 11, 693–702 (2008).

29. Jha, A., Diehl, B., Scott, C., McEvoy, A. W. & Nachev, P. Reversed Procrastination by Focal Disruption of Medial Frontal Cortex. Curr. Biol. 26, 2893–2898 (2016).

30. Heitz, R. P. & Schall, J. D. Neural Mechanisms of Speed-Accuracy Tradeoff. Neuron 76, 1–7 (2013).

31. Boldt, A., Blundell, C. & De Martino, B. Confidence modulates exploration and exploitation in value-based learning. Neurosci. Conscious. 2019, 1–12 (2019).

32. Peters, M. A. K. et al. Perceptual confidence neglects decision-incongruent evidence in the brain. Nat. Hum. Behav. 1, 1–8 (2017).

33. Zylberberg, A., Roelfsema, P. R. & Sigman, M. Variance misperception explains illusions of confidence in simple perceptual decisions. Conscious. Cogn. 27, 246–253 (2014).

34. Maniscalco, B., Peters, M. a K. & Lau, H. Heuristic use of perceptual evidence leads to dissociation between performance and metacognitive sensitivity. Atten. Percept. Psychophys. 78, 923–37 (2016).

35. Van Den Berg, R. et al. A common mechanism underlies changes of mind about decisions and confidence. Elife 5, 1–21 (2016).

36. Kohl, C., Spieser, L., Forster, B., Bestmann, S. & Yarrow, K. Centroparietal activity mirrors the decision variable when tracking biased and time-varying sensory evidence. Cogn. Psychol. 122, 101321 (2020).

37. Peirce, J. et al. PsychoPy2: Experiments in behavior made easy. Behav. Res. Methods 51, 195–203 (2019).

38. McKinney, W. Data Structures for Statistical Computing in Python. Proc. 9th Python Sci. Conf. 51–55 (2010).

39. Delorme, A. & Makeig, S. EEGLAB: An open source toolbox for analysis of single-trial EEG dynamics including independent component analysis. J. Neurosci. Methods 134, 9–21 (2004).

40. Bates, D., Mächler, M., Bolker, B. M. & Walker, S. C. Fitting linear mixed-effects models using lme4. J. Stat. Softw. 67, 251–264 (2015).

